# Species-specific variation in mitochondrial genome tandem repeat polymorphisms in hares (*Lepus* spp., Lagomorpha, Leporidae) provides insight into their evolution

**DOI:** 10.1101/2024.03.04.583303

**Authors:** Riikka Tapanainen, Koit Aasumets, Zsófia Fekete, Steffi Goffart, Eric Dufour, Jaakko L. O. Pohjoismäki

## Abstract

The non-coding regions of the mitochondrial DNAs (mtDNAs) of hares, rabbits, and pika (Lagomorpha) contain short (∼20 bp) and long (130–160 bp) tandem repeats, absent in related mammalian orders. In the presented study, we provide functional annotation for mountain hare (*Lepus timidus*) and brown hare (*L. europaeus*) mtDNA non-coding regions, together with a species- and population-level analysis of tandem repeat variation. Mountain hare short tandem repeats (SRs) as well as other analyzed hare species consist of two conserved 10 bp motifs, with only brown hares exhibiting a single, more variable motif. Long tandem repeats (LRs) also differ in sequence and copy number between species. Mountain hares have four to seven LRs, median value five, while brown hares exhibit five to nine LRs, median value six. Interestingly, introgressed mountain hare mtDNA in brown hares obtained an intermediate LR length distribution, with median copy number being the same as with conspecific brown hare mtDNA. In contrast, transfer of brown hare mtDNA into cultured mtDNA-less mountain hare cells maintained the original LR number, whereas the reciprocal transfer caused copy number instability, suggesting that cellular environment rather than the nuclear genomic background plays a role in the LR maintenance. Due to their dynamic nature and separation from other known conserved sequence elements on the non-coding region of hare mitochondrial genomes, the tandem repeat elements are unlikely to have regulatory roles but are likely to represent selfish genetic elements.

## 1 Introduction

Mitochondria are essential cellular organelles best known for the production of ATP through a process called oxidative phosphorylation (OXPHOS) (Spinelli and Haigis, 2018). They are also a central hub for most metabolic pathways and exhibit specialized functions such as the biosynthesis of iron-sulfur clusters, steroids, fatty acid oxidation, calcium homeostasis, programmed cell death, and more. Due to their origin as a free-living prokaryote, mitochondria have their own small circular DNA genome (mitochondrial DNA, mtDNA), which has undergone reductive evolution associated with gene transfer to the nucleus (Adams and Palmer, 2003). In animals, mtDNA is typically less than 20 kb in size and contains genes for 13 polypeptides of the OXPHOS complexes as well as two mitochondrial ribosomal RNAs and 22 tRNAs required for their translation (Boore, 1999). The two strands of the mtDNA have very different gene density, with most of the genes being encoded on the so-called heavy strand (H-strand). The designation stems from its high guanine content, giving it a higher mass compared to the complementary light-strand (L-strand), as visualized in alkaline CsCl density gradient centrifugation (Kasamatsu et al., 1971). Besides the highly compact coding region, the mitochondrial genome also has a non-coding region (NCR), sometimes called the control region due to the presence of control elements required for DNA transcription and replication. The length and organization of the NCR is very variable across the animal kingdom, contributing most to the mtDNA length variation between the species (Boore, 1999).

In vertebrates, the mtDNA non-coding region contains interesting features, such as the displacement loop (D-loop), which is generated by replication initiating from the main replication origin (OriH: Origin of Heavy strand replication) and terminating prematurely at the “termination associated sequence” (TAS) (Jemt et al., 2015). As a result, a triple-stranded DNA structure is formed, including a 600–700 nt long nascent H-strand fragment, the 7S DNA, which remains hybridized to its L-strand template. While the human and mouse mitochondrial genomes, probably the most studied mtDNAs, show considerable economy in their NCR organization, its size can be variable in other vertebrates due to tandem repeat arrays (Lunt et al., 1998). Of these, short tandem repeats of around 10 bp are an universal feature for the insect mitochondrial genomes (Solignac et al., 1986), but are also present in vertebrate species, including mammals (Savolainen et al., 2000). Longer, 100-200 bp repeats are found in certain species of fish (Wang et al., 2007), birds (Mundy and Helbig, 2004; Omote et al., 2013; Wang et al., 2015) and some mammals, such as rabbits and hares (Casane et al., 1997). The repeat number can vary from two (Wang et al., 2007) to more than one hundred (Hoelzel, 1993) per mtDNA molecule. As tandem repeats can be easily lost during the sequencing and assembly of short sequence reads, such as the ones obtained from Illumina sequencing platforms, it is likely that they are an overlooked feature of mitochondrial genome variation across different taxa. For example, an analysis involving long read sequencing data from vertebrate genome assemblies discovered tandem repeats and gene duplications from several species (Formenti et al., 2021). Interestingly, the presence of tandem repeats was not systematic, but they seem to have been lost and obtained independently in different evolutionary lineages.

The tandem repeats on the NCR of the European rabbit (*Oryctolagus cuniculus*) mitochondrial genome were noted already in the 1980s (Ennafaa et al., 1987) and later work showed that these repeats are widespread also in other mitochondrial genomes of Lagomorpha (Casane et al., 1997). There are two types of repeated motifs in rabbits, 20 bp short repeats (SR) and 153 bp long repeats (LR). SR arrays are expanded or contracted relatively dynamically, with the copy number of the units varying between three to 19. Consequently, the SR arrays can occur in heteroplasmy, i.e. mitochondrial DNAs with different repeat numbers coexisting in the same cell or tissue. An elegant experiment where rabbit SRs were cloned and maintained in a bacterial plasmid demonstrated that slipped-strand mispairing during replication is the main mechanism to explain the dynamic state of the repeat array (Pfeuty et al., 2001). The experiments demonstrated that SR insertions are more common than deletions in arrays with less than 10 copies, with the opposite being true for longer tracts, resulting in the SR lengths to oscillate around an optimal value.

Similar to the SRs, the LRs also have a high mutation rate (10^-2^ per animal per generation), resulting in NCR length variation between generations as well as within individuals, manifesting as heteroplasmy as well as mosaicism (Casane et al., 1997). In rabbits, LR tend to be present as arrays of five repeats with shorter and longer variants being rarer. While the mtDNA haplotype has no influence of the array length, oddly, some organs such as gonads maintain longer arrays than other tissues (Casane et al., 1997). Due to their length, the expansion or deletion of LRs through replication slippage-mispairing is more complicated than for the SRs. In the nucleus, the copy number changes of similar long satellite repeats, including expansions and contractions of rDNA arrays (Kobayashi and Ganley, 2005), have been proposed to occur through break-induced recombination (Thakur et al., 2021). Interestingly, a similar copy-choice recombination mechanism involving strand invasion of the parental molecule by a free 3′-end of a newly synthetized DNA strand at regions of short sequence homologies has been suggested as a mechanism for mtDNA deletion formation in humans (Persson et al., 2019; Phillips et al., 2017). Non-allelic recombination or end joining could in principle also explain the observed LR length variation.

LR tandem repeats have been found in all Lagomorphs from pikas (*Ochotona* spp.) to hares (*Lepus* spp., *Sylvilagus* spp.) (Casane et al., 1997; Dufresne et al., 1996). All LRs have a 20 bp strictly conserved sequence, which has been proposed to be a binding site for a regulatory factor, such as TFAM (Dufresne et al., 1996). In fact, the rabbit LR has been suggested to contain both L-strand (LSP) and H-strand (HSP) promoters for mtDNA transcription (Dufresne et al., 1996). If this is the case, variation in LR length could influence primer synthesis by MTRPOL at these promoters (Kuhl et al., 2016), thereby modulating also the replication initiation.

Our group has been interested in hybridization between the mountain hare (*Lepus timidus*) and brown hare (*L. europaeus*) at the northern contact zone of these species in Finland. Despite being separated by three million years of evolution (Ferreira et al., 2021), the hybrids are fertile, resulting in gene flow across the species barrier (Levanen et al., 2018a; Levanen et al., 2018b). This gene flow is biased towards the brown hare, which seems to benefit from the hybridization by obtaining locally adapted alleles from the mountain hare (Pohjoismaki et al., 2021). Furthermore, the hybridization results in frequent introgression of mountain hare mtDNA into the brown hare population, encompassing up to fifth of the individuals in certain regions in Finland (Levanen et al., 2018a). When sequencing the mitochondrial genomes of our mountain hare and brown hare cell lines (Gaertner et al., 2023), we recently noted a species difference in the length of LRs in the mtDNA non-coding region. A population sampling of 151 mountain hares and 148 brown hares with species-specific mtDNA confirmed that brown hares maintain more LR copies (median = 6) than mountain hares (median = 5), with no correlation with their geographic origin. Interestingly, when introgressed into brown hares, the mountain hare mtDNA presented frequent heteroplasmy and gained “brown-hare like” LRs (median = 6 copies). To further test how the nuclear background can influence LRs, we generated cybrids of mountain hare and brown hare fibroblasts with nuclear DNA from one species and mtDNA from another. The repeat region length changed in three cybrid cell lines out of six, regardless of their genetic background, with the new LR variant maintained in heteroplasmy. In one case, the LR number of brown hare mtDNA was increased from six to seven in cells with mountain hare nucleus, a state which is very rare in the natural population of mountain hares. We conclude that while the nuclear genetic background certainly plays a role in the LR maintenance, environmental factors and differences in cell biology between species also contribute to the observed variation. We also discuss the potential relation of the NCR repeat elements to the regulation of mtDNA.

## 2 Materials and Methods

### 2.1 Sampling and DNA isolation

The mountain hare (*Lepus timidus*) and brown hare (*L. europaeus*) specimens presented here are from a larger biobank of 1,202 hare samples collected in 2012–2016 across Finland (Levanen et al., 2018a; Levanen et al., 2018b). The origin and generation of the four mountain hare (LT1, LT4, LT5, LT6) and four brown hare (LE1, LE2, LE3, LE4) fibroblast cell lines has been described elsewhere (Gaertner et al., 2023). Most samples are from hunted animals, with a subset from specimens found dead from the wild. The sampling had minimal impact on the local populations and posed no threat on the habitats. The hunting followed the regional hunting seasons and legislation (Metsästyslaki [Hunting law] 1993/615/5§). The sampling adhered to the ARRIVE guidelines and no ethical assessment was required. All sampling occurred in Finland and did not involve International Trade in Endangered Species of Wild Fauna and Flora (CITES) or other export of specimens, as defined by the Convention on Biological Diversity (CBD). The DNA was isolated from frozen ear muscle biopsies using the peqGOLD blood & tissue DNA mini kit (VWR) and following the protocol provided by the manufacturer.

### 2.2 Mitochondrial DNA sequencing

The primers used to amplify the hare mtDNA were as follows:

Le93F: TTGTTTTGTAGCAAGTTTACACATGC

Le184R: GCTTAATACCTGCTCCTCTTGATCTA

Le1580F: TTAAACCCATAGTTGGCCTAAAAGC

Le1635R: TTGAGCTTTAACGCTTTCTTAATTGA

Le3045F: AGGCGTATTATTTATCCTAGCAACCT

Le3175R: CCTCATAAGAAATGGTCTGTGCGA

Le3921F: CCCCCTAATCTTTTCCATCATCCTAT

Le4482R: TCATCCTATATGGGCAATTGAGGAAT

Le4689F: AGGCTTTATTCCAAAGTGAATTATTATTCA

Le5417R: AGGCTCCAAATAAAAGGTAGAGAGTT

Le6696F: ATACCGTCTCATCAATAGGCTCCTTC

Le6756R: ATAAAGATTATTACTATTACAGCGGTTAGA

Le8603F: AGCCTATATCTACATGATAATACTTAATGA

Le8698R: CGGATAAGGCCCCGGTAAGTGG

Le10552F: TTGAAGCAACACTAATCCCTACACTA

Le10613R: TCGTTCTGTTTGATTACCTCATCGT

Le11301F: ACCATTAACCTTCTAGGAGAGCTTCT

Le11807R: AGGATAATGATTGAGACGGCTATTGA

Le12407F: GTCTAATCCTAGCTGCTACAGGTAAG

Le12791R: GAGCATAAAAAGAGTATAGCTTTGAA

Le14204F: ATTGTTAACCACTCTCTAATCGACCT

Le14514R: CCAATGTTTCAGGTTTCTAGGTAAGT

Lt16056F: TGGGGTATGCTTGGACTCAAC

Le16119R: TCGTCTACAATAAGTGCACCGG

In total, 12 separate reactions were prepared to cover the mitochondria genome:

1. Lt16056F + Le184R: 1871 bp

2. Le93F + Le1635R: 1543 bp

3. Le1580F + Le3175R: 1596 bp

4. Le3045F + Le4482R: 1438 bp

5. Le3921F + Le5417R: 1497 bp

6. Le4689F + Le6756R: 2068 bp

7. Le6696F + Le8698R: 2003 bp

8. Le8603F + Le10613R: 2011 bp

9. Le10552F + Le11807R: 1256 bp

10. Le11301F + Le12791R: 1491 bp

11. Le12407F + Le14514R: 2108 bp

12. Le14204F + Le16119R: 1916 bp

(Expected fragment size based on the published *Lepus europaeus* mtDNA sequence from Sweden [NC_004028.1]).

The fragments were amplified from total DNA preparations using a PCR program with a 1 min 94 °C denaturing step, followed by 35 cycles of 94 °C for 15 s, 56 °C for 15 s and 72 °C for 2 min and a final 3 min elongation step at 72 °C. The obtained products were gel purified using the GeneJET gel extraction kit (Thermo Fisher Scientific™) and sent for sequencing using Illumina MiniSeq™ at the Genome Center of Eastern Finland.

The sequence of the non-coding region-containing PCR fragment (Lt16056F + Lt184R) was further validated by Sanger sequencing, applying also the following additional sequencing primer:

Le101F: TATAAATTCCTGCCAAACCCCAAAAA

### 2.3 Mitochondrial DNA assembly and annotation

Mitochondrial DNA was assembled using the MitoZ pipeline(Meng et al., 2019) and the best assembly was selected by comparing the results of the pipeline’s outputs with different kmer options. The final assembly was done using the un options --clade Chordata –fastq_read_length 150, --requiring_taxa Chordata --genetic_code 2 --kmers_megahit 21 29 39 59 79 99 119 141. The pipeline included the tools fastp (Chen et al., 2018) for cleaning the raw data, MEGAHIT (Li et al., 2015) for assembly, after which sequences were filtered using HMMER (Wheeler and Eddy, 2013) to ensure the correct taxa and the completeness of protein-coding genes. Annotation was performed using TBLASTN (Gertz et al., 2006), GeneWise (Birney et al., 2004) and MiTFi (Juhling et al., 2012). The annotation of the non-coding region (NCR) as well as the illustration of the mitochondrial genome was done with Geneious® 10.2.6 (Biomatters. Available from https://www.geneious.com). The functional loci on the NCR were identified based on the similarity with the human (NC_012920) and mouse (FJ374652) NCR sequences.

### 2.4 Genotyping of the long repeat (LR) region

The LR length genotyping of the hare samples was performed by PCR using the following primers that bind the flanking regions of the repeat run (Fig. 1A, B):

**Fig. 1.**
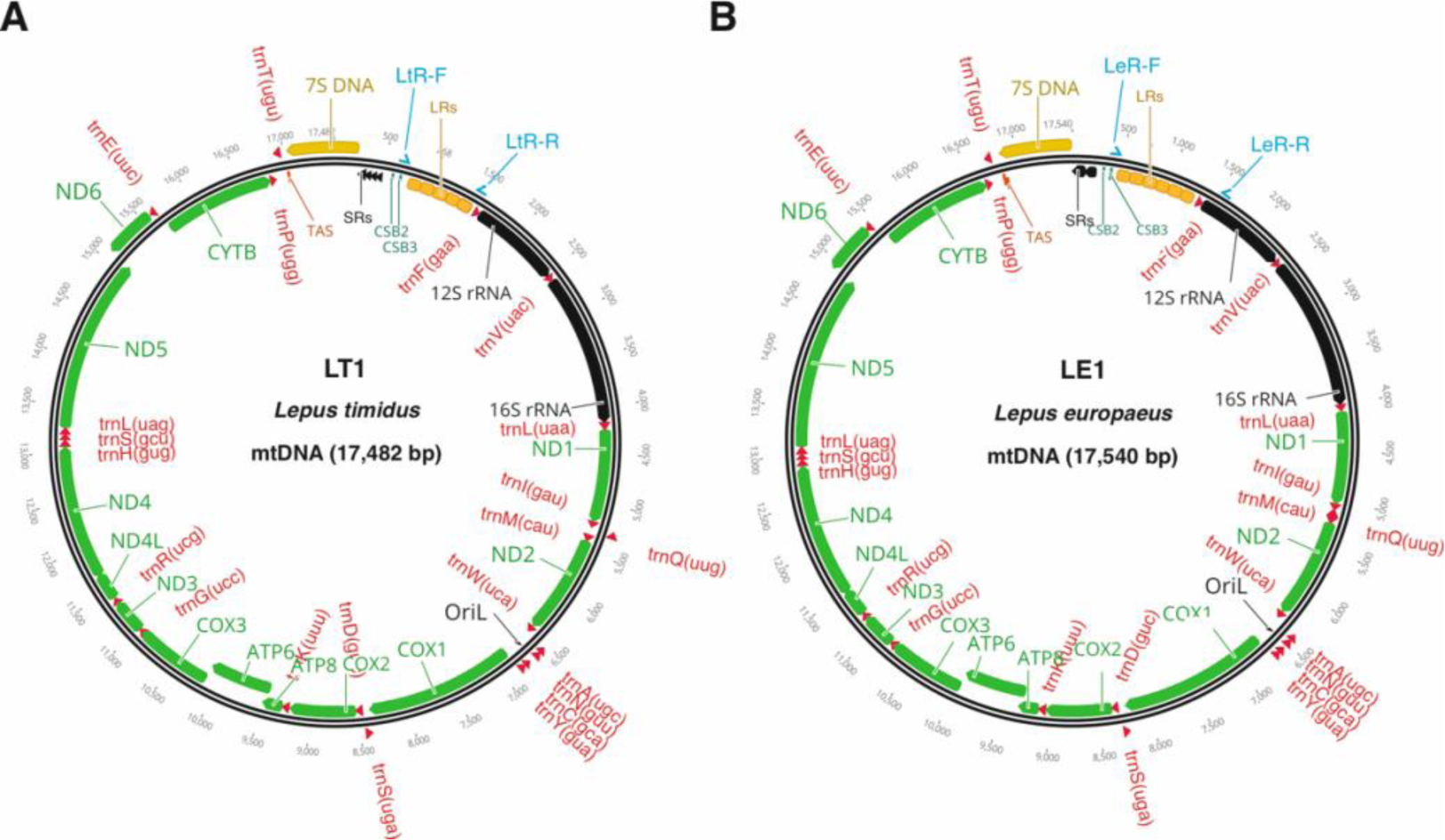
Schematic illustration and gene maps of hare mitochondrial genomes. (A) The mitochondrial DNA from LT1 mountain hare (*Lepus timidus*) and (B) LE1 brown hare (*L. europaeus*) fibroblasts. The binding sites for the long tandem repeat (LR) genotyping primers (LR-F, LR-R) are indicated. See Fig. 2 for the details of the non-coding region.

*Lepus timidus* mtDNA

LtLR-F: AGAACCGTGACATAGCACTTACTTTC

LtLR-R: TAACATATTGGTGTTAGAATGTTTTTAGT

*L. europaeus* mtDNA

LeLR-F: TATAAATTCCTGCCAAACCCCAAAAA

LeLR-R: GCTTAATACCTGCTCCTCTTGATCTA

The used PCR program had an initial denaturation step of 94 °C for 2 min, followed by 35 cycles of 94 °C for 20 s, 59 °C for 20 s and 72 °C for 90 s, with final elongation at 72 °C for 5 min, using AccuStart II™ PCR SuperMix (Quantabio). The PCR products were separated over a 1 % TAE agarose gel.

### 2.5 Analyses of the SRs

Core SR sequences and their flanking regions of the Finnish *Lepus timidus* and *Lepus europaeus* were manually retrieved from the whole mitochondrial genome sequences. All possible core dimers (1 for LE type, 4 for LT) were then used as query sequence for a basic nucleotide BLAST search. Results were filtered to only include mitochondrial genomes sequences from defined geographic isolates (LE query) or from one genome per *Lepus* species (LT queries). For LT queries; *Lepus tibetanus*: MN539746.1, *L. tolai*: MN539744.1, *L. arcticus*: NC_044769.1, *L. sinensis*: NC_025316.1, *L. coreanus*: NC_024259.1, *L. granatensis*: NC_024042.1, *L. othus*: KJ397608.1, *L. corsicanus*: KJ397606.1, *L. capensis*: NC_015841.1 and *L. yarkandensis*: MN539747.1 mitochondrial genomes were selected. *L. townsendii* was excluded due to poor quality sequencing of its SR region. For the LE query, *L. europaeus* Poland 3: KY211034.1, Poland 2: KY211033.1, Poland 1: KY211032.1, Germany 2: KY211031.1, Germany 1: KY211030.1, Greece 4: KY211029.1, Greece 3: KY211028.1, Greece 1: KY211026.1, Cyprus 4: KY211025.1, Cyprus 3: KY211024.1, Cyprus 2: KY211023.1, Cyprus 1: KY211022.1, and Turkey 1: KY211021.1 were selected. LE Greece 2 isolate was excluded as its SR region was identical to that of Greece 3 isolate. Rabbit (*Oryctolagus cuniculus*) reference genome mtDNA (NC_001913.1) was used for comparison. The proportion of mutated core repeats was compared using Fisher exact test. Because of its unique SR features, *L. yarkandensis* was excluded from the statistical analyses. Sequence uncertainties in the repeats of *L. otus* (1) and *L. corsicanus* (3) were all considered as mutated for the Fisher exact test, but the mutation frequency was nevertheless significantly higher in the LE type of SRs. To visualize the frequency of mutation in function of the repeat location, the repeat positions were normalized to the number of repeats in the SR leading to an x-axis ranging from 0 (“beginning” of the SR region) to 1 (“end” of the SR region).

### 2.6 Generation of cybrid cell lines

Cybrid cells were generated using the chemical enucleation method (Bayona-Bafaluy et al., 2003). In brief, first recipient mtDNA-less ρ0 cells from immortalized LT1, LT4, LT6, LE1, LE2 and LE3 fibroblast cell lines were generated by culturing them in the presence of 100 μM ddC, until no mtDNA was detected on a Southern blot. The ρ0 cells were maintained in high glucose (4.5 g/l) DMEM, supplemented with 10 % FBS and 50 μg/ml uridine. The nucleus of the mtDNA donor cells was removed with Actinomycin D, with the effective concentration determined separately for each cell line. The optimal concentration for most cell lines was 1 μg/ml for 18 h, except for LT1 (1 μg/ml for 15 h) and LE2 (2 μg/ml for 28 h). The cybrid fusions were conducted in both directions (same cell line either a recipient or a donor of mtDNA) for the following cell line pairs: LE1♂ x LT4♀, LE2♀ x LT1♂ and LE3♀ x LT6♂. For the control cybrids with mtDNA from the same species but a different cell line than the nucleus, we used LE1♂(nucleus) x LE2♀(mtDNA) and LT4♀ (mtDNA) x LT6♂ (nucleus) combinations. The fusion was performed by growing the mtDNA donor cells on 6-well plates and treating them with Actinomycin D. After replacing the treatment medium with fresh DMEM containing 10 % FBS 1 million ρ0 cells were added to the well for 3 h. The cells were then washed 3 × with fresh DMEM before addition of 45 % polyethylene glycol (MW 1450 g/mol) for 60 s. The cells were then washed again 3 × with DMEM + 10 % DMSO and once with DMEM. Finally, the cells were given DMEM + 10% FBS + Penicillin/Streptomycin. Unlike the ρ0, the cybrid cells were maintained without added uridine to select for functional mitochondrial DNA. The cells were grown to confluency, at which stage half of the cells in a well were collected for genotyping. DNA was extracted with Quick-DNA Miniprep Kit (Zymo Research).

As the donor and recipient cells differed by their sex, PCR-RFLP of *ZFX* and *ZFY* loci (Fontanesi et al., 2008) was used to genotype the cybrid nucleus and *CYTB* PCR-RFLP their mtDNA (Melo-Ferreira et al., 2005). Each original cybrid pool was genotyped and promising looking pools were subcloned on 96-wells. These clonal cell lines were rechecked for the expected nuclear and mtDNA haplotypes, obtaining the final cybrid cell lines.

### 2.7 Agarose gel electrophoresis and Southern hybridization

Mitochondrial DNA from LE and LT parental and cybrid cell lines was isolated using phenol:chloroform extraction and ethanol precipitation. 2 µg of uncut mitochondrial DNA were separated over a 0.4 % agarose gel in 1x TBE buffer at 25 V overnight at room temperature. To facilitate the transfer of mtDNA onto nylon membrane by capillary transfer, the gel was subjected to acid depurination followed by denaturation to create suitable hybridization targets. The transferred DNA was crosslinked by baking 80 °C for 2 h. The Southern blot was probed using (α-32P)-dCTP labelled PCR probe spanning the hare mtDNA nucleotides 16840-88 or 17077-88 at 65 ℃ for overnight. Signals were captured on a phosphor screen and detected using a phosphorimager (Fujifilm FLA-3000).

### 2.8 Statistical tests

The RNAfold web server (http://rna.tbi.univie.ac.at/cgi-bin/RNAWebSuite/RNAfold.cgi) for predicting secondary structures of single stranded RNA or DNA sequences was used to screen for hairpin formation in the mitochondrial non-coding region and the QGRS mapper (Kikin et al., 2006) to identify putative Quadruplex forming G-Rich Sequences.

To test whether there was a geographical pattern in the LR length variation among Finnish hares, a spatial autocorrelation analysis was used applying the Moran’s I test with sf and spdep packages in R (https://CRAN.R-project.org/).

## 3 Results

### 3.1 The structure of the mitochondrial DNA non-coding region in hares

We amplified the mitochondrial genomes from four mountain hare (LT1, LT4, LT5, LT6) and four brown hare (LE1, LE2, LE3, LE4) cell lines as 12 overlapping PCR products, which were subsequently sequenced using Illumina Miniseq™. When assembling the sequences, we noted that the obtained sequence for the NCR was shorter than expected from the PCR product lengths. Closer inspection revealed that this was because the short sequence reads (2 × 150 bp) over the LR region were assembled as a single repeat element due to their redundancy. The PCR products containing NCR elements were therefore reanalyzed using Sanger sequencing to clarify LR copy number and finalize the assemblies (Table 1, Fig. 1). Interestingly, our mountain hare as well as brown hare mitochondrial genomes are smaller (Table 1) than the previously published mountain hare mtDNA from a Finnish specimen (NC_024040.1) (17,755 bp) or the brown hare mtDNA from Sweden (NC_004028.1). The size difference is explained by the length of the non-coding region (NCR), where our LT cell lines have five and LE cell lines six long tandem repeats (LRs) compared to seven in both previously published mitochondrial genomes.

**Table 1.**
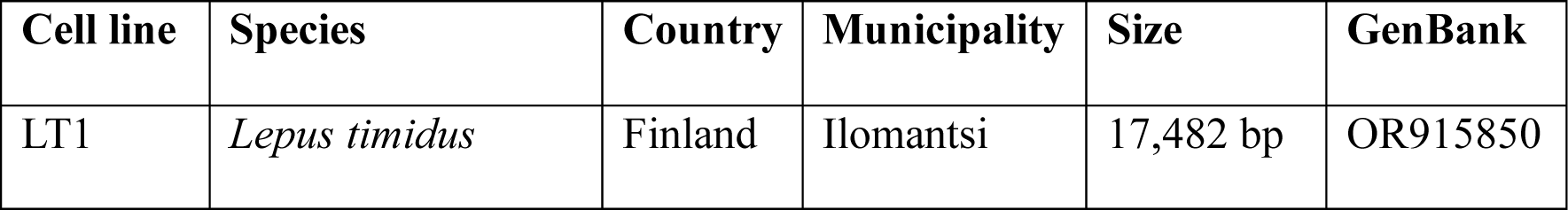
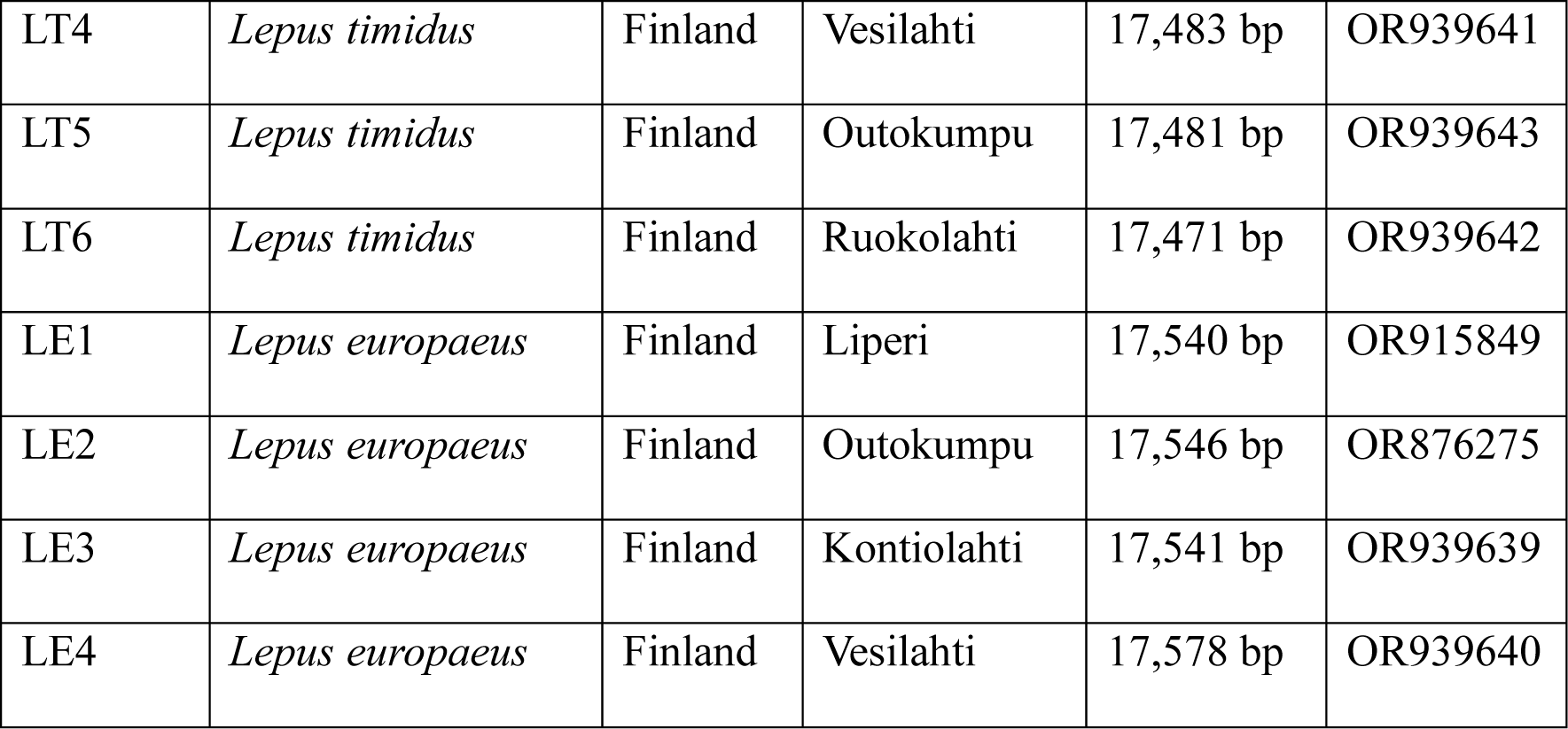
The mitochondrial genomes from mountain hare (LT) and brown hare (LE) cell lines sequenced in this study.

The length of the NCR is 2,038 bp in LT1 (Fig. 2A) and 2,102 bp in LE1 (Fig. 2B). It contains recognizable features such as a rather conserved region between the termination associated sequence (TAS) and the cluster of short tandem repeats (SR), which by length and location could correspond the D-loop region containing the 7S DNA. The SRs and the LRs are separated by a variably long sequence containing the conserved sequence blocks (CSBs) 2 and 3 thought to be relevant for the control of mtDNA gene expression and replication priming (Pham et al., 2006). However, if replication is primed at or near the CSBs, the primer 3′-ends are relatively far away from the known 5′-ends of DNA, considered to correspond the replication origin and start of the 7S DNA (Pohjoismaki et al., 2018). In fact, Southern blot analysis of this region can only detect the 7S DNA when using a probe that extends downstream of the SRs (Fig. 2C). Also, the size of the 7S does not differ between the cell lines despite the difference in the length of the NCR-region. Of note, we could confirm that Lagomorphs do not seem to have any CSB1(Casane et al., 1997; Dufresne et al., 1996). The CSB2 differs by a couple of indels between the species, while the CSB3 seems highly conserved (Fig. 3A).

**Fig. 2.**
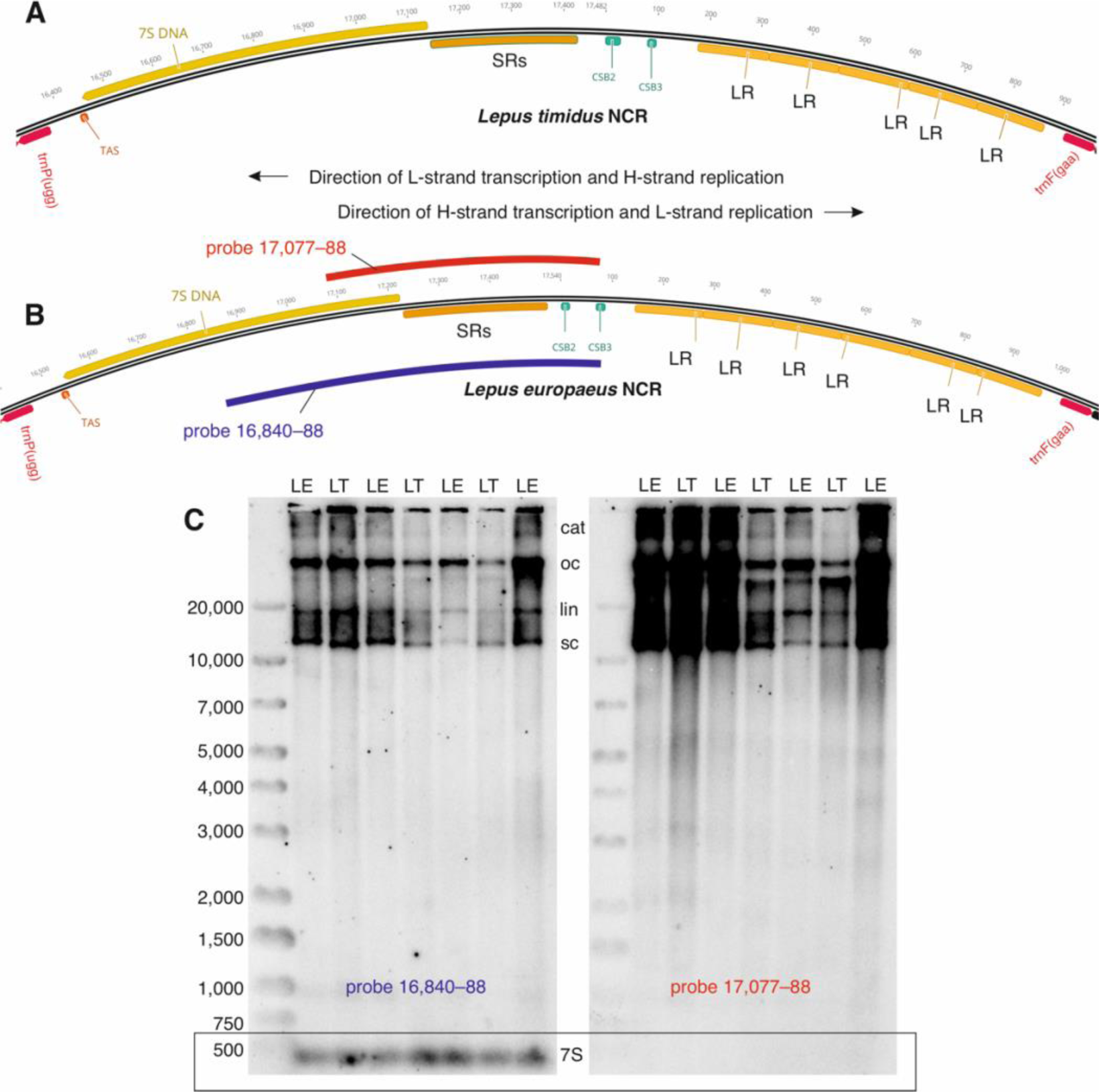
Sequence features of hare mitochondrial DNA non-coding region (NCR). (A) The NCR of LT1 cells and (B) LE1 cells. The direction of transcription and replication, as well as probe locations for the Southern blot are indicated. Note that the H-strand replication is the leading-strand, initiated first and primed by the L-strand transcript. Key: trnP = tRNA-proline; TAS = termination associated sequence; SR = short tandem repeat; CSB2 and 3 = conserved sequence block 2 and 3; LR = long tandem repeat; trnaF = tRNA-phenylalanine. (C) A Southern blot of uncut mtDNA from mountain hare (LT) and brown hare cells (LE), probed for the region spanning nts 16,840–88 (left panel) or 17,077–88 (right panel). See the probe locations in (B). The right panel is overexposed to demonstrate the absence of 7S signal. Key: cat = catenated mtDNA forms, oc = open circles; lin = linear mtDNA; sc = supercoiled circles; 7S = 7S DNA.

**Fig. 3.**
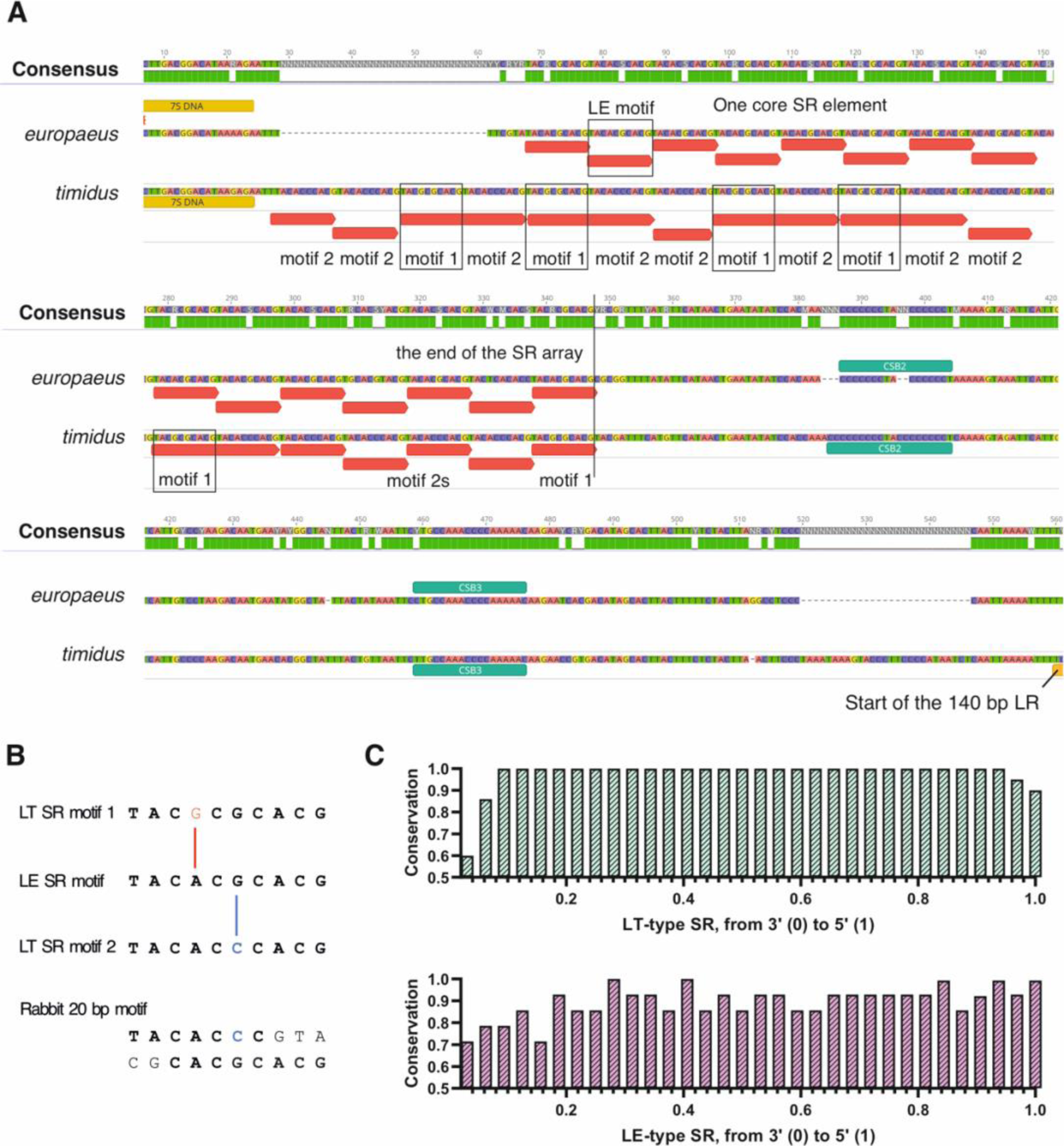
Comparison of the sequence differences between the brown hare (*L. europaeus*) and mountain hare (*L. timidus*) mtDNA non-coding regions between the assumed start of the 7S DNA and the long tandem repeats. (A) The short tandem repeats (SRs) form a continuously repeating array of a single motif in brown hare, while the mountain hares show random mixtures of two different repeat motifs. Repeating sequences and their direction are marked with red bars. Mountain hare CSB2 also has more Cs flanking the central TA-pair, whereas the CSB3 sequence is identical between the species. Note both SNP and indel variation between the two species. (B) The 10 bp conserved motif sequences in the hare SRs. The LE type is only seen in brown hares whereas LT types are present in all hares analyzed here. Note the G-A transition (red) and G-C transversion (blue) between the three core sequences. The 20 bp rabbit SR motif is shown in two parts as comparison. Identical sequences in bold. Note the similarity of the two rabbit sequence halves with the brown hare repeat motif. (C) Sequence conservation of the core SR sequence motifs in hares. The LT type shows remarkable conservation across species, whereas the LE type shows more variation within one species.

### 3.2 Hare SR elements consist of 10 bp core motifs

At first glance, the mountain hare (LT) SRs appear as a 20 bp repeat element similar in length to the rabbit’s SRs (Fig. 3A). However, these are not consistent, and part of the SR arrays showed repetition of only half of the sequence. On closer inspection, the element consists of two 10 bp motifs, TAC<u>G</u>C<u>G</u>CACG, and TAC<u>A</u>C<u>C</u>CACG, which differ by the two underlined nucleotides. In contrast, the brown hare (LE) SRs contain only one type of 10 bp core motif (TAC<u>A</u>C<u>G</u>CACG), which differs from both LT motifs by one of the variable nucleotides (Fig. 3B). Interestingly, the 20 bp rabbit SR motif contains the 7 first nucleotides of the brown hare (LE) SR motif spliced together with the last 8 nucleotides of LT SR motif1, separated by an unique 5 bp (GTACG) spacer sequence (Fig. 3B).

While the LE type is present only in brown hares, the two LT types occur in all other hare species that we analyzed (Table 2), with the two core elements alternating variably in the SR arrays. Curiously, also the cape hare (*Lepus capensis*), a close relative of brown hare (Ferreira et al., 2021), has LT type SRs. *Lepus yarkandensis* constitute another curiosity, with randomly alternating core 1 and 2 LT type SR motifs being separated by dinucleotides (TT or TC). Overall, the LT type SRs show considerably less variation than the LE types (Fig. 3C, 3.35%; *p* < 0.001, Fisher’s exact test), despite being spread across many species. The copy number of the LT core repeat sequences varied from 32 in our LT cell lines and other *L. timidus, coreanus, granatensis, otus* and *corsicanus* isolates to 16 in *L. tibetanus* (Table 2). The copy number of the LE type, present only in brown hares, ranged from 28 to 32.

**Table 2.** The short repeat core element types and copy number variation among hares. The clade indicates whether the species is closer related to mountain hare (LT) or brown hare (LE). Core element copy numbers represent only the analyzed mitochondrial genomes (see materials and methods) and not a population study. LT+ for *Lepus yarkandensis* indicates a unique dimer insertion in the classical LT type SR (see text).

| Species | Species clade | SR type | Core element copy number |
| --- | --- | --- | --- |
| *Lepus timidus* | LT | LT | 32 |
| *Lepus tibetanus* | LT | LT | 16 |
| *Lepus tolai* | LT | LT | 17 |
| *Lepus arcticus* | LT | LT | 23 |
| *Lepus sinensis* | LT | LT | 28 |
| *Lepus coreanus* | LT | LT | 32 |
| *Lepus granatensis* | LT | LT | 32 |
| *Lepus otus* | LT | LT | 32 |
| *Lepus corsicanus* | LT | LT | 32 |
| *Lepus yarkandensis* | LT | LT+ | 21 |
| *Lepus capensis* | LE | LT | 29 |
| *Lepus europaeus* | LE | LE | 28–32 |

As repetitive sequences are likely to form secondary structures, we were interested to see whether the SR sequences of the two hares can form hairpins when single-stranded. This is particularly interesting as hairpin formation at the second most prominent replication origin on mitochondria, the light-strand origin (OriL) located at the WANCY tRNA cluster, is proposed to be required for L-strand replication priming (Fuste et al., 2010; Wanrooij et al., 2008). A rather strong hairpin was predicted for the H-strand SRs from the mountain hares but not from the brown hares, nor on the L-strand SR sequences of either species (Fig. 4A–C). As for other structural features, the entire non-coding region contained only a single sequence stretch at the 5′ end of the 7S DNA predicted to form a strong guanine quadruplex (G4) (Fig. 4D).

**Fig. 4.**
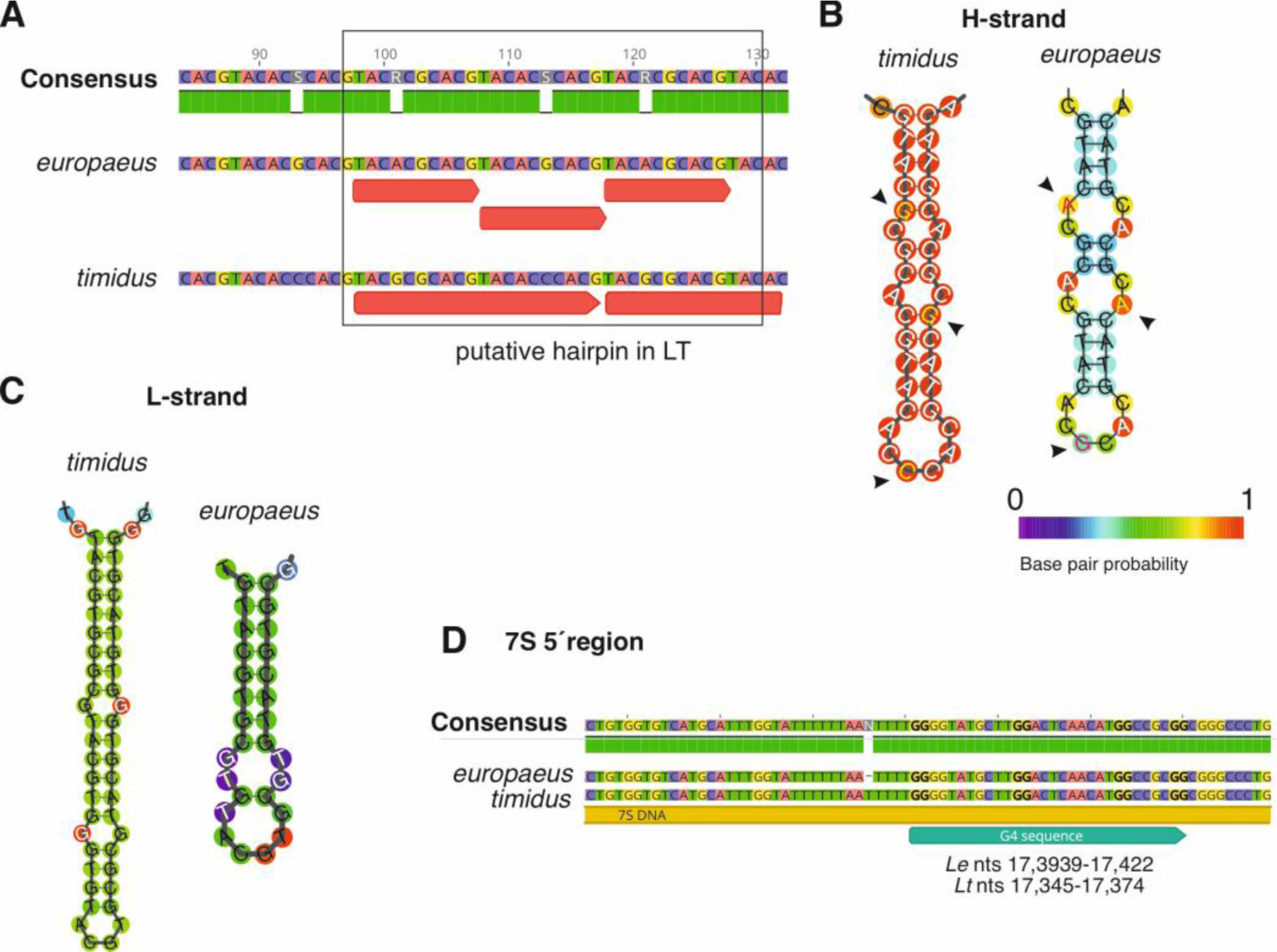
Structural features of the hare mitochondrial non-coding region. (A) Despite the difference in the repeat element length and their overlap, there are only a few SNPs between the short repeat elements (SR) from the brown hare (*europaeus*) and mountain hare (*timidus*). (B) Predicted mountain hare and brown hare SR H-strand secondary structures. Note the longer and stronger hairpin formation in mountain hare. (C) Predicted secondary structures on the SR L-strand. (D) A putative G-quadruplex sequence close to the 5′ end of the 7S DNA (nts 17,078–17,107 in LE1). G-duplets are highlighted.

### 3.3 LR length variation in Finnish hares

The LRs of the two species are similar in length (Fig. 5A), 139 bp in the mountain hare and 140 bp in the brown hare, differing at 21 nucleotide locations and containing the same 20 bp conserved sequence element as previously reported for all Lagomorpha (Casane et al., 1997). As also noted earlier, the LR copy number shows variation between and even within individuals (Fig. 5B, C). Although PCR tends to cause the amplification of short repeat sequences as an artefact, which are evident as a faint ladder of PCR products of varying repeat content, the common repeat haplotype was readily detectable as a strong band in all population samples and systematically confirmed the sequencing results.

**Fig. 5.**
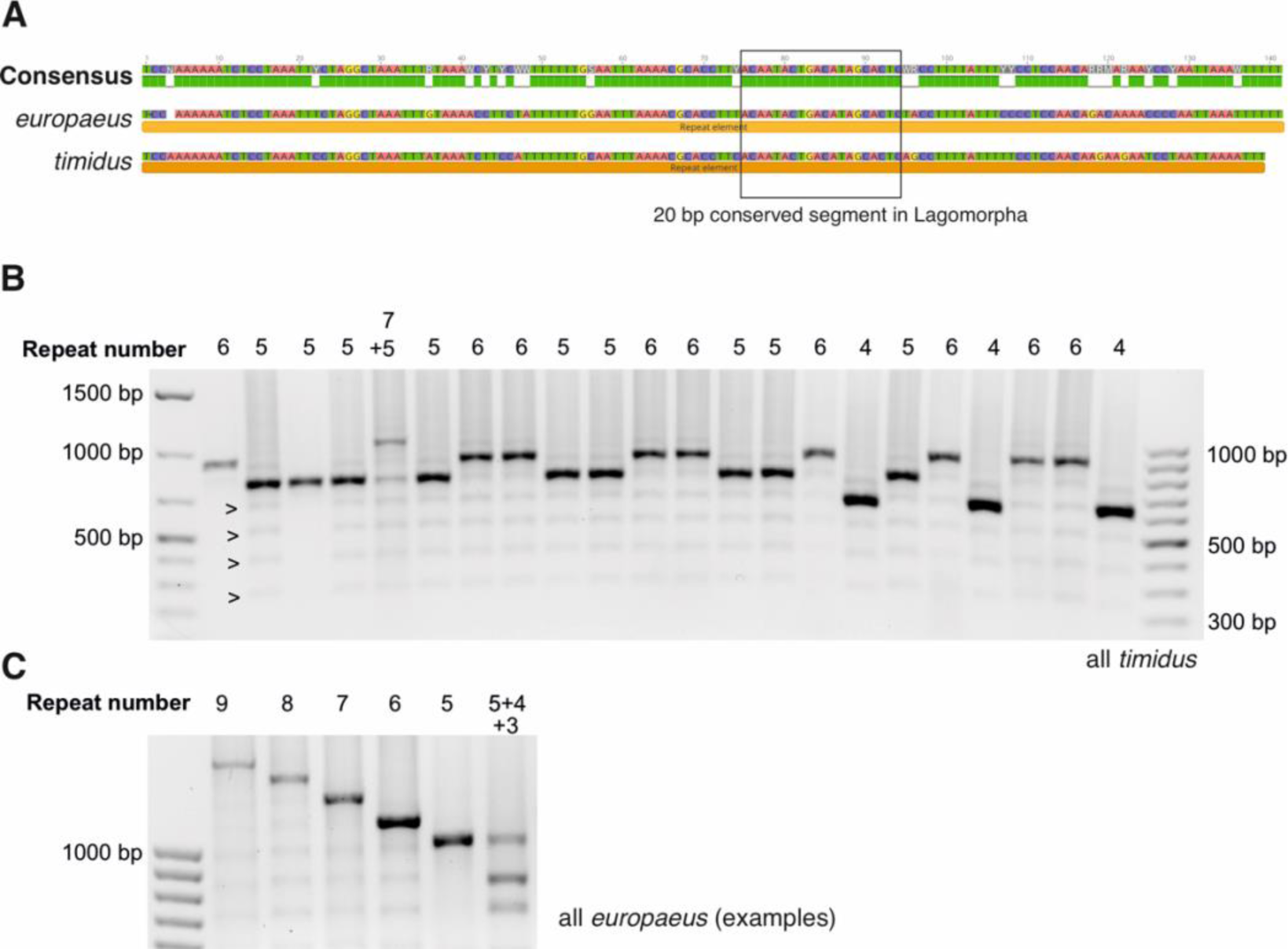
Long tandem repeats (LR) in hares. (A) Sequence comparison of LRs from brown hares and mountain hares, highlighting the 20 bp conserved sequence present in all known Lagomorpha. (B) An example of LR copy number variation in mountain hares. Arrowheads highlight LR ladder generated as a PCR artefact. (C) Examples of LR copy number variants present in brown hares. Size difference compared to (B) due to different PCR primers.

While the mechanism of dynamic length variation in the SRs has been shown to result from replication slippage (Pfeuty et al., 2001), the factors influencing the LR array length are less clear. As they are also known to have species differences (Lunt et al., 1998), we were interested to see whether the LR array length would show population-specific variation. The most common length of LR was five repeats in Finnish mountain hares, with six repeats being the next abundant (Fig. 6A). The shortest observed repeat was four and longest seven, which was represented only by one sample. Curiously, the only previously published mountain hare mtDNA from Finland (NC_024040.1) also has seven repeats. Heteroplasmy was detected in five samples (3 %). No geographic pattern in the LR length variation was observed among the 151 mountain hares across Finland (Moran I statistic standard deviate = 8.09×10^-10^, p-value = 0.5).

**Fig. 6.**
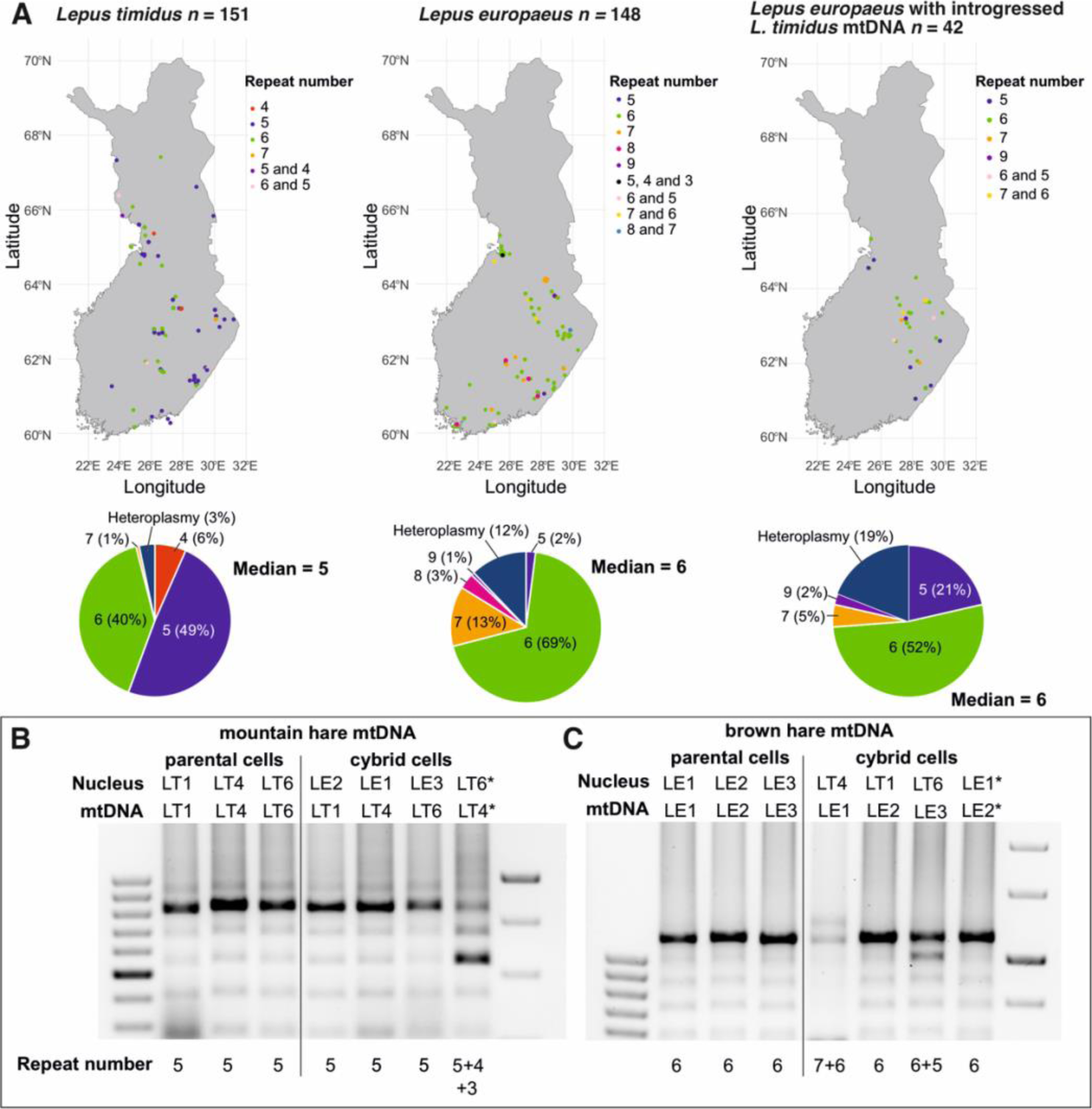
Long tandem repeat (LR) copy number variation in Finnish hares. (A) The geographical distribution of LR copy number variants and their frequencies in Finnish mountain hares (*Lepus timidus*) and brown hares (*L. europaeus*) with conspecific or introgressed mtDNA. (B) LR copy numbers in the original mountain hare fibroblasts and xenomitochondrial cybrids with mtDNA from these cells. (C) The same for the parental and cybrid cells with brown hare mtDNA. LT = *Lepus timidus*; LE = *L. europaeus.* Asterisk indicates control cybrids with nucleus and mtDNA from the same species, but different parental cell lines.

In contrast to mountain hares, the most common LR region length in brown hares was six repeats, with seven and eight repeats being more common than five (Fig. 6A). The longest observed LR region had nine repeats, while shorter than five repeats occurred only in heteroplasmic samples (Fig. 5C). In the heteroplasmic samples, the shortest repeat sequence length was three copies. Overall, brown hares showed more frequent heteroplasmy (12 %, *p* = 0.0045, Fisher exact test). Interestingly, brown hares with introgressed mountain hare mtDNA showed an intermediate LR-region length distribution mtDNA (Fig. 6A). Although they had the same median length of six repeats as brown hares with conspecific mtDNA, five repeats were much more common (21 % vs. 2 %). Also, a notable proportion (19 %) of the introgressed samples were heteroplasmic, although the difference to the other brown hares was not significant (*p* = 0.3077, Fisher exact test). Again, no correlation between the LR length and geography was observed in brown hares with or without introgressed mtDNA. Only a few mountain hares with brown hare mtDNA were present in our collection (Levanen et al., 2018a) and therefore were not included in the analysis.

### 3.4 Transfer of mtDNA between cells causes instability in the LR copy number

Our LT and LE fibroblasts had five and six LR repeats, respectively. If the repeat length is an autonomous property of mitochondria, we expect it to remain stable despite the cohabitation with a different nuclear genome. If it is controlled by the nuclear genome, we expect it to be rapidly converted to the copy number corresponding to the nuclear donor. While no change in the LR copy number was observed when mtDNA was transferred from mountain hare to brown hare cells (Fig. 6B), the six copies LR array length of brown hare mtDNA increased to seven in LT4, remained the same in LT1 and decreased to five in LT6 nuclear background (Fig. 6C). Curiously, novel LR copy numbers of four and three appeared also in control cybrids with LT4 mtDNA in LT6 nuclear background (Fig. 6B), with no change observed in the brown hare control cybrids (Fig. 6C). In all cases, the new LR array haplotypes existed in heteroplasmy with the original haplotype.

## 4 Discussion

When sequencing the mitochondrial DNAs as a part of the basic characterization of our hare fibroblast cell lines (Gaertner et al., 2023), we noted a discrepancy between the expected genome size and the assemblies obtained from short-read Illumina data. Closer inspection revealed that this was because of misassembly of the part of the non-coding region with known long tandem repeats (LRs). We corrected the assemblies with Sanger sequencing and provide here the fully annotated mitochondrial genomes for four mountain hares and four brown hares from Finland (Table 1, Figs. 1–2). Although tandem repeats are a known elements of the non-coding region in many vertebrate species, including lagomorphs, they remain an overlooked aspects of mitochondrial genome variation, largely due to their absence in humans and other well-studied model organisms. In addition, and as in our case initially, they are likely to be lost when assembling mitochondrial genome sequencing data obtained using short-read sequencing platforms. For this reason, a lot of hare mitochondrial genome assemblies available in GenBank have only one LR-element and many of the SR-array sequences might have issues as well. It is likely that the same problem applies to the mitochondrial genome assemblies from other organisms, meaning that the frequency and diversity of such elements in vertebrates is probably underestimated. Recent development of long-molecule sequencing technologies can help to complete our understanding of the repeat element variation, especially as the full-length mitochondrial genomes can be assembled reliably from whole-genome sequencing data (Uliano-Silva et al., 2023).

Not much is known about the function and biological significance of the repeat elements in the non-coding region of mitochondrial genomes. Initial studies proposed that the Lagomorph LRs are involved in the regulation of transcription and replication initiation (Casane et al., 1997; Dufresne et al., 1996). However, there are other conserved regulatory elements on the NCR (Fig. 2), which are known to function in e.g. mtDNA replication priming in other mammals (Pham et al., 2006). Furthermore, the 7S DNA, resulting from prematurely terminated replication, does not contain sequences between LR- and SR-regions (Fig. 2C), indicating that the replication is likely to start at the SR, as suggested by other authors (Melo-Ferreira et al., 2014).

Interestingly, the SR array length can change rather dynamically in rabbits (Casane et al., 1997; Dufresne et al., 1996; Pfeuty et al., 2001) and their copy number is highly variable (from three to 19). In hares, the nature of the repeating sequences made the delimitation of the actual repeat element difficult, as different types of repeats with varying length can be identified (Fig. 3A). Like in rabbits, a 20 bp SR element can be identified in mountain hares. However, this element consists in fact of two separate motifs (Fig. 3B), which can occur also alone as the repeating element in the SR array. In brown hares, the SR arrays consist only of one core 10 bp motif, which differs from both mountain hare motif types by a single nucleotide (Fig. 3B). If only these canonical sequences are counted, their copy number in the different hare species varied between 16 and 32 (Table 2). Considering the evolution of the SRs, it is interesting that the LT and LE core motifs also form the basis of the 20 bp rabbit SR, differing by five interrupting nucleotides, GTACG. It is noteworthy that these can be converted by C>T and A>G transitions to form a LT motif 2 + LE motif 20 bp consensus sequence block (Fig. 3B). It is likely that such transition mutations can occur repeatedly during evolution, while the G>C transversion, separating the LT motif 2 and 3 part of the rabbit motif from the rest (Fig. 3B), is unlikely as a recurrent point mutation. Alternatively, it is possible that the LE motif and LT motif 2 are recombinants of two ancestral repeat types, which are present (in modified state) in rabbits. Overall, the remarkable conservation of the core LT-type SR repeats in hares (*Lepus*) is intriguing (Fig. 3C). This suggests that their origin predates the separation of these species and highlights the unique situation of *L. europaeus* SRs, whose core sequence is not shared by the other *Lepus* species, including its close relative, the cape hare (*L. capensis*) (Ferreira et al., 2021). Nevertheless, the identical length of the SRs core sequences (10 bp) and their similarity in all *Lepus* species (Fig. 3B, only a single nucleotide difference separates the *L. europaeus* core sequence from each of two SR cores found in other *Lepus* species) can be only explained by a common ancestry or mechanism of origin among all species in the genus. The sequence homogeny of the SR runs within a species is quite likely maintained by constant contraction and reamplification of the repeat motifs, resulting in their rapid drift.

Three more observations are worth mentioning concerning the SRs region. (a) In all species and independently of the SR core, the repeats located at the 3′ end and in the center of the SR are the least conserved (Fig. 3C). Similar asymmetry is known to result from the unidirectional gene conversion associated with some mitochondrial mobile elements or with mating type switching in yeast (Richard et al., 2008; Stoddard, 2014), which includes a double-strand break followed by erosion of the ends that generates nucleotide variation after repair. (b) Amongst the 48 core sequence variants in LE-type and LT-type of repeats, no LT-type core variant could be observed in species with LE-type core repeats and *vice versa*, even though the SR core sequences show very strong homology,. This is suprising, as it suggests that a unique SR motif amplification event has taken place specifically in the brown hare evolutionary lineage. Sequencing mitochondrial SRs from more *Lepus* species and isolates would be needed to allow drawing further hypotheses from this observation. (c) One species, *L. yarkandensis*, presents a unique SR motive constituted of LT motif 1 + LT motif 2 + a two-nucleotide (TT or TC) spacer. Similar short footprint signatures have been associated to the excision of some transposons (Skipper et al., 2013) as well as non-homologous end joining repair mechanisms (Yant and Kay, 2003).

The biological significance or functional roles of the SRs remain enigmatic. It is plausible that the SRs could form secondary structures when being single-stranded, which could promote priming by MTRPOL, as occurs during the initiation of L-strand replication at OriL (Fuste et al., 2010). Interestingly, hairpin formation is predicted to be stronger for the H-strand (Fig. 4), which – following the OriL priming mechanism – would indicate a L-strand origin. Similarly, the reported variation in the SR copy number (Casane et al., 1997; Dufresne et al., 1996) and their proneness to mutate through strand-slippage (Pfeuty et al., 2001), are counterintuitive features for important genomic functions such as the regulatory control of DNA-replication.

Like the short repeats, the long repeats contain a core (20 bp) sequence segment (Fig. 4A), which is conserved in all lagomorphs (Casane et al., 1997; Dufresne et al., 1996). Coincidentally, this is the same length as the duplex SR motifs, although the sequence bears no resemblance to them and has no obvious sequence features such as palindromes. If it represents a regulatory element capable of binding protein factors such as TFAM, as proposed (Dufresne et al., 1996), the existence of multiple such sites on mtDNA is somewhat reminiscent of the initiator titration model of bacterial replication initiation (Hansen et al., 1991; Ogawa et al., 2002). In *E. coli* for example, the initiation factor DnaA is titrated away from the replication origin *oriC* binding sites by providing competing recognition sites dispersed elsewhere on the genome. Replication of the genome causes the duplication of these sites, binding more DnaA and thereby reducing its availability to initiate new rounds of replication. The obvious problem for this mechanism to operate in the regulation of hare mtDNA replication initiation is the variability in the copy number of the LRs (Fig. 4B, C), where having three to nine copies of competitive binding sites per genome would effectively negate any accurate regulation, although this could be balanced by a difference in mtDNA copy number and TFAM molecules. Unlike the dispersed DnaA sites, the LRs are clustered together and situated so that their doubling would occur only at the end of the replication. Thus, they are unlikely to prevent re-initiation before termination. Still, they could control replication by titrating TFAM in function of the number of mtDNA copies (and therefore SRs repeats) already present in the mitochondria. However, as with the SRs, if these elements have such an important role in genome regulation, why would they be present in some evolutionary lineages but not in others? For example, rodents (Rodentia) – a sister order of Lagomorpha – as well as most other mammals do not have such repetitive sequence arrays on their mtDNAs.

The species difference in the LR length distribution between brown hare and mountain hare (Fig. 6A) is interesting, as it tells that their variation between individuals is not random. In fact, similar differences in mtDNA tandem repeat lengths are known from other closely related species (Hernández et al., 2004; Lunt et al., 1998; Mundy and Helbig, 2004; Omote et al., 2013). As the species difference is fundamentally a genetic difference, one might expect that the LR copy number would then also be governed by the nuclear genome. However, the fact that LR copy number can vary between the tissues of the same individual in rabbits and that certain tissues are more prone to have longer arrays than others (Casane et al., 1997), suggests a more complicated mechanism, where the nuclear genetic background is only one variable. As different tissues have different metabolic requirements, manifesting as differences in mitochondrial activity and mtDNA maintenance strategies (Herbers et al., 2019), it is plausible that the internal environment in mitochondria also indirectly effects the LR-region length through factors such as temperature, oxidative stress and replication/repair protein availability. In fact, we previously noted that mountain hare fibroblasts maintain higher mitochondrial membrane potential than brown hare cells (Gaertner et al., 2023) and have also other differences in their oxidative metabolism. In this case, the energetic state of the cells can be influenced by variables such as metabolic adaptations, tissue type, ageing and stress, causing also the observed variation in the LR copy numbers. If the LRs are generated or lost as a result of strand-slippage during replication, as seems to be the case with the SRs (Pfeuty et al., 2001), changes in the mitochondrial internal environment could increase the probability of such events, resulting in LR copy number change. Similarly, factors such as mitochondrial temperature (Chretien et al., 2018), could favor certain repeat lengths over others and stabilize the LR copy number variation. Overall, it is curious that the SR core sequence shows less conservation (Fig. 3C) and the LR arrays show more length variation in brown hares than in mountain hares.

Interestingly, our experiments with the cybrids showed mixed results (Fig. 6B). First, the effect on LR copy number was asymmetric and only presented by cells with mountain hare nucleus, regardless of the mtDNA origin. Also, the change in the LR array length in was not consistent. While in one cybrid line the brown hare mtDNA lost a LR copy, obtaining the common copy number of five for a mountain hare mtDNA, in another the copy number was increased to seven, a LR array length not presented by any of the parental cell lines. In contrast, shorter than parental mountain hare LR arrays appeared in the control cybrid with LT6 nucleus and LT4 mtDNA. Although the sample size is small, it is interesting that the observed effect is consistent with the frequent introgression of mountain hare mtDNA into brown hare population but not *vice versa* (Levanen et al., 2018a; Thulin and Tegelström, 2002), suggesting that mtDNA maintenance is perhaps more permissive or resilient in cells with brown hare nucleus. Alternatively, the mountain hare cells could be more sensitive to the stress caused by the process of generating cybrids. In all cases, the new haplotype appeared in heteroplasmy with the old, showing that they arise as a spontaneous rearrangement of the existing LRs and increase in abundancy through drift or selection.

## 5 Conclusions

Although being a decades old discovery, repeat elements on the non-coding region remain an overlooked feature of Lagomorph mitochondrial genomes. We recognized conserved 10 bp core sequences in the short repeats, with two different motifs present in most hare species and one in brown hares. Interestingly, SR core element copies showed sequence variation in brown hares, but not in other hare species. We also found that while the long repeat arrays are dynamic, their presentation is not random, as the different species show different length distributions. Considering their absence from most mammalian species, as well as the lack of a clear biological role in mtDNA maintenance and expression, it is likely that both repeat types have arisen and are maintained as selfish genetic elements. It is unlikely that these elements are mobile, and they are certainly too small to encode functional gene products, indicating that their reproduction has to be related to the mechanism of mtDNA replication. Why such elements have not been lost from the otherwise compact mitochondrial genomes and how species-specific copy number variation is maintained, warrants further research.

## 6#Acknowledgements

Dr Seppo Helisalmi and Dr Joose Raivo from the Gene diagnostics lab at Genome Center of Eastern Finland are thanked for providing the mtDNA sequencing service. We thank Ms Anita Kervinen for her valuable laboratory assistance in preparing the mtDNA PCRs for the sequencing. Funding: This research was supported by the Academy of Finland (grant no. 329264 for ED and JLOP). The funder had no role in study design; sampling, analysis and interpretation of data; in the writing of the report; and in the decision to submit the article for publication.

